# Force-exerting lateral protrusions in fibroblastic cell contraction

**DOI:** 10.1101/711507

**Authors:** Abinash Padhi, Karanpreet Singh, Janusz Franco-Barraza, Daniel J. Marston, Edna Cukierman, Klaus M. Hahn, Rakesh K. Kapania, Amrinder S. Nain

## Abstract

Aligned extracellular matrix fibers enable fibroblasts to undergo myofibroblastic activation and lead to elongated cell morphology. The fibroblasts in turn contract to cause alignment of the extracellular matrix. This feedback process is critical in pathological occurrences such as desmoplasia and is not well understood. Using engineered fiber networks that serve as force sensors, we identify lateral protrusions with specific functions and morphology that are induced by elongated fibroblastic cells and which apply extracellular fiber-deflecting contractile forces. Lateral projections, named twines, produce twine bridges upon interacting with neighboring parallel fibers. These mature into “perpendicular lateral protrusions” (PLPs) that enable cells to spread laterally and effectively contract. Using quantitative microscopy, we show that the twines originate from the stratification of cyclic actin waves traversing the entire length of the cell. The primary twines swing freely in 3D and engage neighboring extracellular fibers. Once engaged, a lamellum extends from the primary twine and forms a second twine, which also engages with the neighboring fiber. As the lamellum fills in the space between the two twines, a sheet-like PLP is formed to contract effectively. By controlling the geometry of extracellular networks we confirm that anisotropic fibrous environments enable PLP formation, and these force-generating PLPs are oriented perpendicular to the parent cell body. PLP formation kinetics indicated mechanisms analogous to other/known actin-based structures. Our identification of force-exerting PLPs in anisotropic fibrous environments suggests an explanation for cancer-associated desmoplastic expansion at single-cell resolution, providing possible new clinical intervention opportunities.

## INTRODUCTION

During acute wound healing, myofibroblastic activation of fibroblastic cells imparts inward forces and contracts the collagen-rich extracellular matrix (ECM). Upon wound resolution, this becomes the acellular scar. Myofibroblastic cell contraction is pivotal to many physiological processes (e.g., developmental, acute wound healing), but is also key to pathological instances like chronic inflammation and fibrosis-related diseases, including microenvironmental desmoplasia in solid cancers^1,2^. It is thus not surprising that cancers have been described as chronic wounds for which desmoplasia, the contractile fibrous-like tumor-microenvironment, plays a pivotal role^3,4^. In solid epithelial tumors, desmoplasia expands in a manner that simulates chronic wound extension^5^. In fact, in extreme cases such as pancreatic ductal adenocarcinomas (PDAC), desmoplasia can expand to encompass the majority of the tumor mass^6^. Interestingly, just as scarred tissue often presents with aligned collagen signatures, aligned ECM fibers were described as “tumor-associated collagen” or TAC signatures, hallmarks of detrimental patient outcomes^7–10^. Importantly, inflammatory and myofibroblastic cancer-associated fibroblasts (CAFs) are known to be the local activated cells responsible for the production and remodeling of the topographically aligned, anisotropic, desmoplastic ECM. Of note, we previously demonstrated that anisotropic ECMs generated by CAFs can activate naïve fibroblastic cells into CAFs^11^, suggesting that aligned ECMs generated by CAFs could drive desmoplastic expansion. Despite extensive data regarding the role of aligned ECM in polarized migration^8,12–16^, very little is known about ECM-dependent desmoplastic expansion. It is commonly understood that in aligned ECMs, cell elongation and polarity are maintained due to minimal probing in lateral directions^15^. In such scenarios, it is not understood how cells stretch onto multiple-fibers, and how lateral protrusions contract inwards to aid in desmoplastic remodeling and expansion. This study identifies and characterizes a new type of contractile, lateral fibroblast protrusion through which aligned ECM triggers desmoplastic expansion.

Lateral protrusions or spikes are well-documented in multiple cell types (growth cones, mesenchymal, fibrosarcoma, hepatocytes, and conjunctival cells)^17,18^ and occur in varying sizes and morphologies to serve unique purposes.^18,19^ These structures have been implicated in the proteolytic degradation of the 3D interstitial ECM by cancer cells^18^, and by leukocytes in probing the vascular membrane for permissive sites, which eventually allow vascular extravasation ^20^. Leukocytes experiencing shear flow ‘roll,’ via transient tethering, using receptor-ligand bonds.^21,22^ The tethering interactions are facilitated by microvilli shown to be 80nm in diameter and varying in length from 350nm to 1μm or longer^23,24^. Similarly, lateral protrusions formed in melanoma cells have been documented to provide traction in a 3D environment in the absence of adhesion.^25^ It was recently shown that spindle-shaped cells in 3D gel matrices maintain cell polarity and directed migration by forming limited numbers of lateral protrusions,^26^ which are attributed to restriction in α_5_β_1_-integrin to the leading edge ^15,27^, and activating RhoA laterally to inhibit Rac1-induced protrusions.^27–29^ Overall, the utility of lateral protrusions remains poorly understood, as classic 2D culturing methods limit the formation of lateral protrusions, while 3D gels lack the homogeneity and topographic patterning needed to study the role of fiber geometry on protrusion frequency, morphology, and dynamics. To partially remedy this, we recently used orthogonal arrangement of fibers of mismatched fiber diameters to constrain cell migration along large diameter fibers while studying lateral protrusions of various shapes and sizes. We observed these formed in an integrin-dependent manner on small diameter fibers^30,31^. Here, using aligned and suspended fiber nanonets that also act as force sensors^32,33^, we report the identification of force exerting side protrusions termed ‘perpendicular lateral protrusions (PLPs)’. We quantitatively show that anisotropic ECMs promote production of PLPs to enable fibroblastic cell contraction, explaining the single cell/ECM physical characteristics of desmoplastic expansion.

## RESULTS

### Twines emerge from actin ruffles and engage neighboring fibers

On suspended and aligned fibers, we observed that elongated cells formed filamentous lateral protrusions (primary *twines*) that swung freely in 3D until they effectively attached to neighboring ECM fibers. Subsequently, we observed initiation and formation of *secondary* twines that transitioned to formation of a twine-bridge, composed of primary and secondary twines, oriented perpendicular to cell body. We observed a sheet-like structure formed to fill the gap between the twines to form “perpendicular lateral protrusions” (PLPs, **Figure 1**, and **Movie M1**). The PLPs were capable of exerting forces that enabled cells to spread in the transverse direction and deflect the neighboring fiber, suggestive of cellular contraction perpendicular to main cell body axis.

**FIGURE 1.**
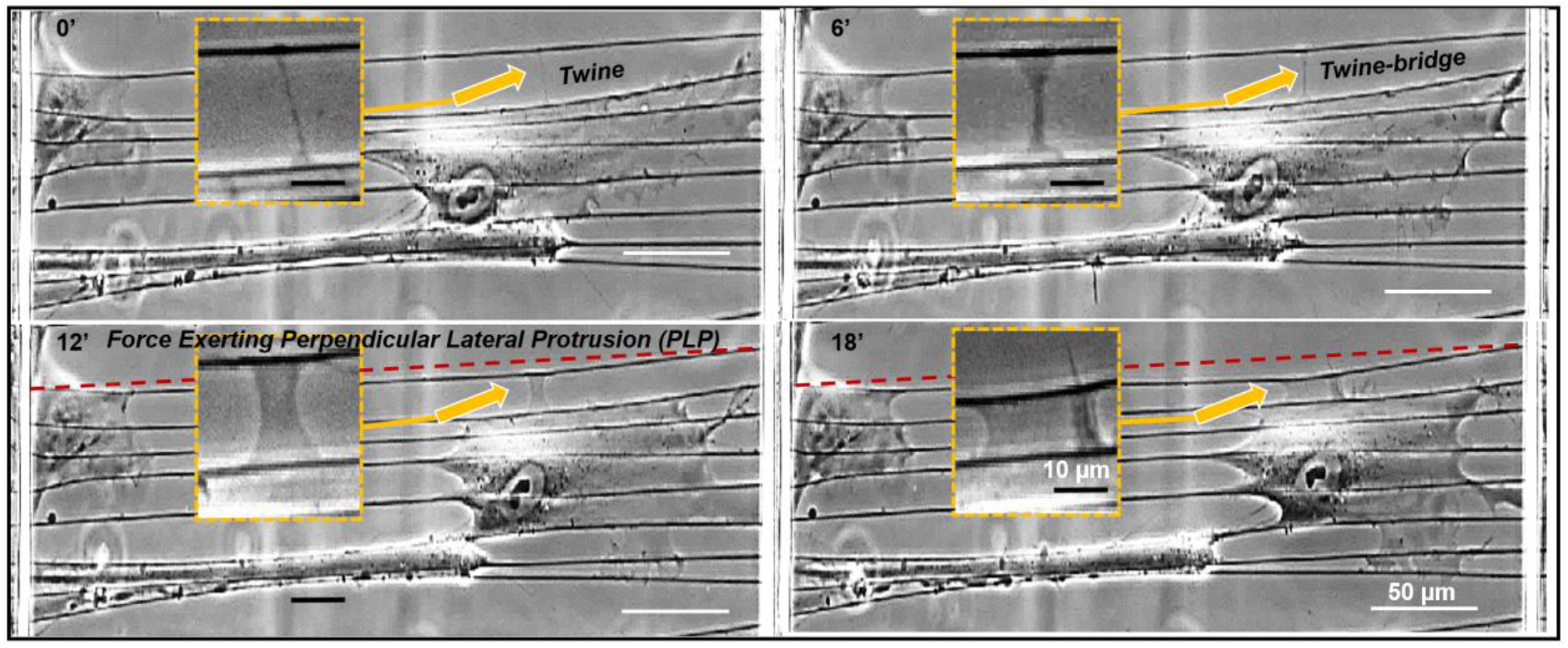
Transition from primary twines to perpendicular lateral protrusions (PLPs). Insets show magnified images of the sections indicated by arrows. The dashed red line shows the original, undeflected position of a fiber neighboring the cell. Timelapse images of single cell in elongated shape spreading laterally through force exerting PLPs on neighboring fibers. A single twine contacts a neighboring fiber (upper left; 0’), leading to production of a second twine. The space between them is filled in to form a twine bridge (upper right; 6’). Twines mature to become capable of exerting inward contractile forces at 12 minutes and proceed to mature and deflect further by 18 minutes (bottom images; 12’ and 18’, compare neighboring fiber deflection vs. red doted line, marking the initial neighboring fiber position).

To quantitatively describe the formation of PLPs, we first inquired if lateral protrusions could form in the elongated cells, and asked if they exhibited a characteristic morphology. For this we used the previously described non-electrospinning Spinneret based Tunable Engineered Parameters (STEP^34–36^) method to fabricate anisotropic/aligned fibers. These could also serve as force sensing nanonets, as we described previously^32,37^. We followed, in real-time, the protrusions of different cell lines to examine which if any behavior were unique to fibroblastic cells (immortalized murine (C2C12, embryonic fibroblast (MEF), and well-established NIH-3T3) and human mesenchymal stem cells (hMSCs) as well as Hela cells) (**Supplementary figure S1** and **Movies M2-M5**). All cell types acquired an elongated shape, as expected, while continuously forming twines (**Figure 1**). The twines were located lateral to the long axis of the cell and found to originate from membrane ruffles that resembled previously described cyclic actin waves^38^. The membrane ruffles spiraled about the fiber axis (**Figure 2A** and **Movie M6**) and advanced at 6.58 μm/min (n=30), similar to the actin polymerization rates reported in the literature (7.20-8.73 μm/min).^39^ The ruffles extended beyond the main cell body and stratified into denser independent twines (**Figure 2B** and **Movie M7**) presumably due to folding-over and buckling.^40^ The twines were found to grow in length and swing freely (in 3D) similarly to reported angular rotations of filopodia,^41–44^ which enabled engagement to neighboring fibers. The attachment to neighboring fibers occurred rapidly following contact between the twine and the neighboring fiber (**Figure 2C** and **Movie M8**).

**FIGURE 2.**
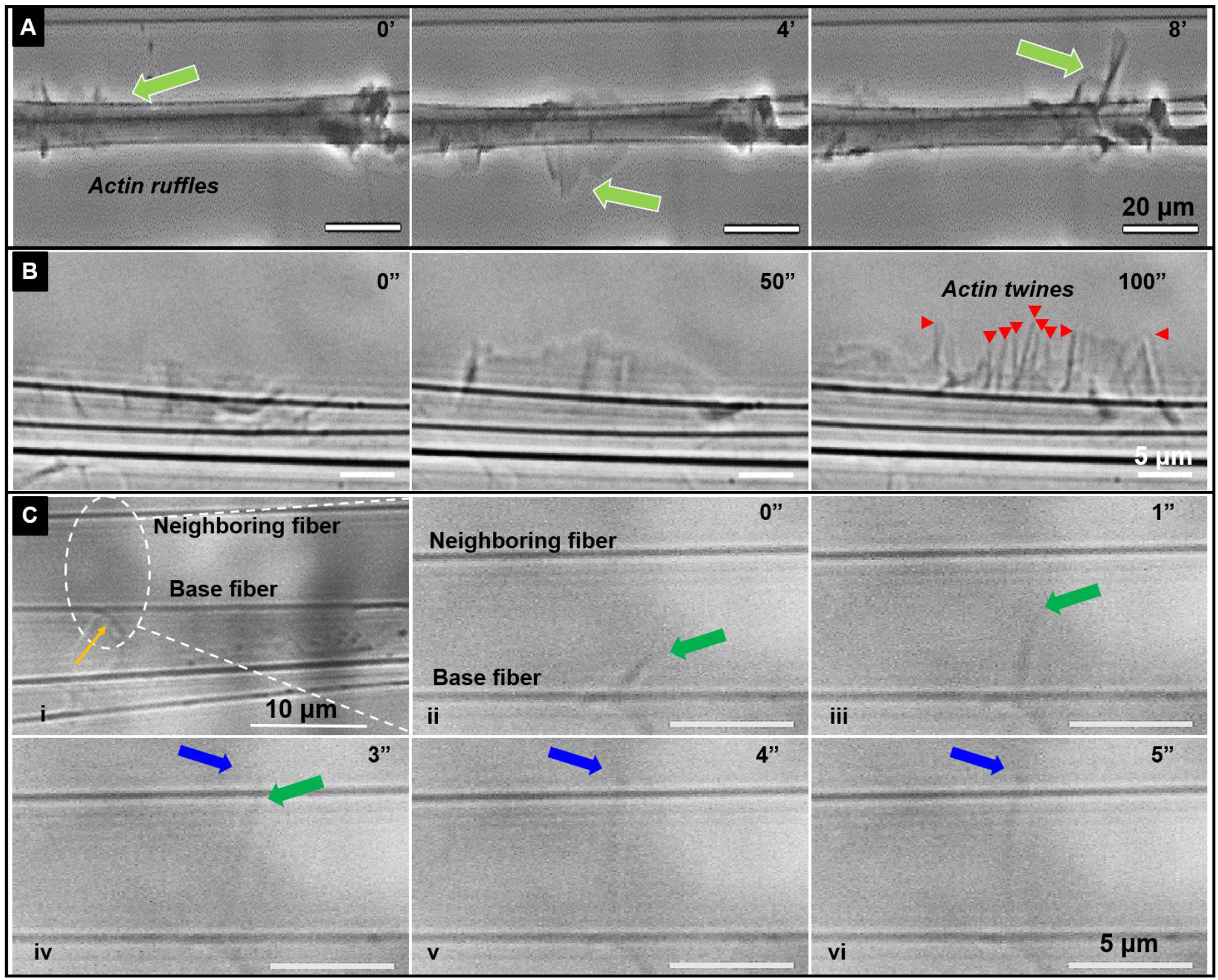
In anisotropic substrates, twines emerge from actin ruffles and engage with neighboring fibers. Time-lapse images showing (A) Actin wave (green arrow) travelling along the length of the cell body (Movie M6). (B) Ruffles from the wave develop into striated structures that evolve into nano-filamentous nascent ‘twines’ shown by red arrowheads (Movie M7). (C) i. Image of a cell with a primary twine pointed by the yellow arrow. (ii-vi) Time-lapse images of area indicated by dashed oval demonstrating that twine engagement with neighboring fiber takes place in ~2 seconds of intial primary twine contacting a neighboring fiber ((iv-vi), Movie M8). Green arrow shows growth of primary twine from cell body and blue arrow shows twine growth after engagement with neighboring fiber i.e. overhanging part of twine.

### Discreet steps in the formation of perpendicular lateral protrusions (PLPs)

We characterized the evolution of individual primary twines into PLPs. Using real-time phase microscopy of living cells, we observed the formation of primary twines, secondary twines, and twine-bridges (**Movie M9**). The maturation of these into force-exerting PLPs occurred in seven characteristic steps (**Figure 3A(a,b)**): (i) Cells stretch, on anisotropic extracellular fibers (‘base fibers’), and achieve an elongated morphology parallel to the fiber orientation. Step (ii) The base fibers serve as anchors enabling the cell to probe for neighboring fibers by extending rigid (see below) “probing twines” lateral protrusions that lunge outwards and sway from their base to encounter and engage with the neighboring fiber (**Figure 2c**). Step (iii) Cell extends a triangular lamellum anchored between the primary twine and the base fiber. Step (iv) From the lamellum, one or more ‘secondary twines’ are formed similar to folding and buckling driven stratification of membrane ruffle described in Figure 2. Step (v) Secondary twines are defined and reach towards the neighboring fiber. When at least one makes contact and adheres to the neighboring fiber, which has a primary twine attached to it, the two twines act as anchors, and the original lamellum begins to fill in the space between the main cell body at the base fiber and the neighboring fiber, thus forming a suspended ‘twine-bridge’ (**Figure 3B**). Step (vi) After the twine bridge completely fills the space between the two twines, the cell body (at the base fiber) and the neighboring fiber (forming an ‘interfiber lamellum’), the twine-bridge widens with a classic hour-glass shape to attain an early PLP (timepoint 6 minutes in **Figure 1**). Step (vii) As the PLP widens/matures, it applies inward contractile forces, thus deflecting the neighboring fiber. The original neighboring fiber can now serve as a new base fiber, thus perpetuating fibroblastic expansion and increased cellular contraction. Surprisingly, we found that these seven steps in PLP formation were not isolated occurrences and importantly were unique to aligned fiber networks (see below).

**FIGURE 3.**
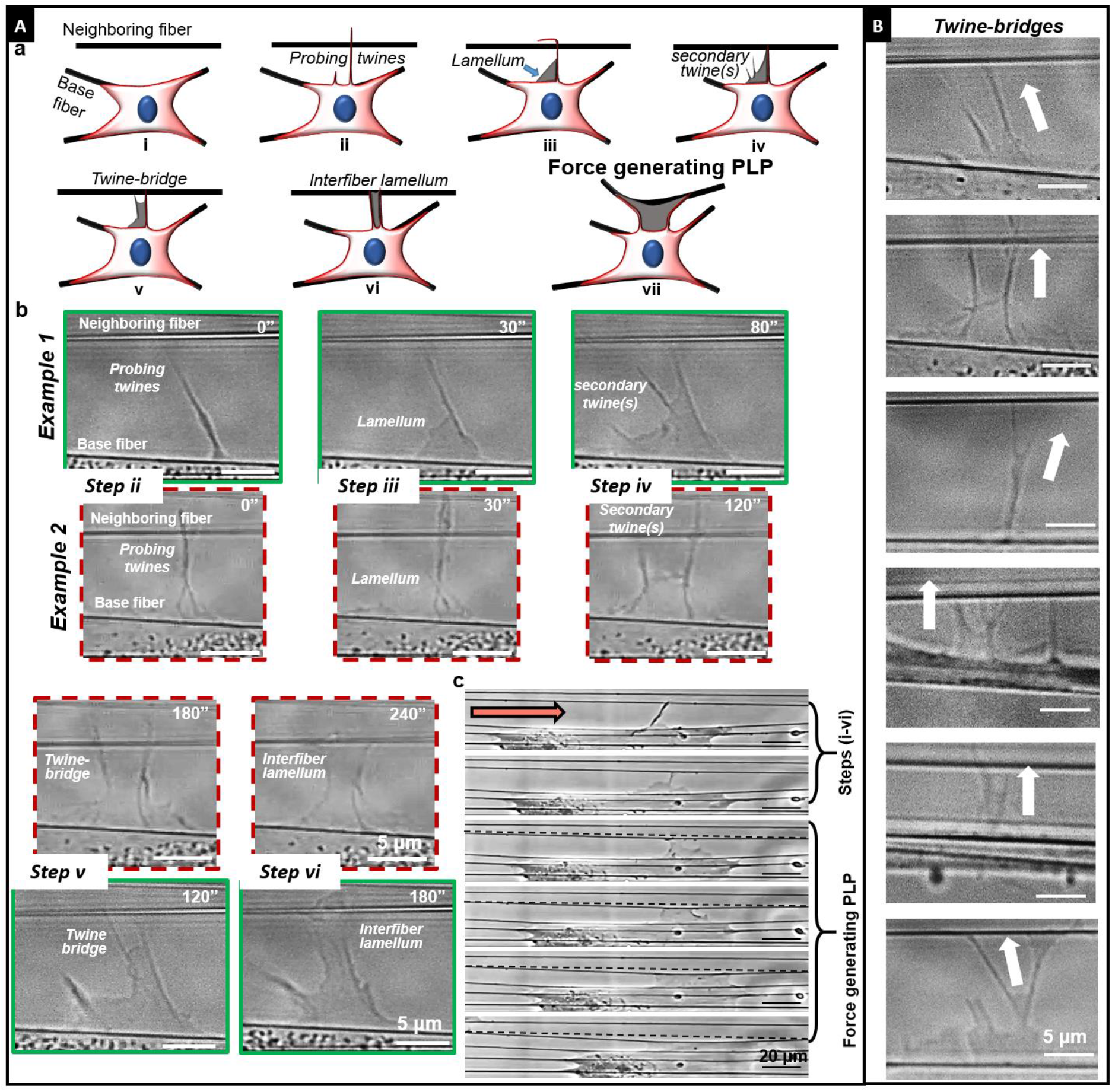
Discreet steps in formaiton of force exerting perpendicular lateral protrusions (PLPs). (A) a) Cartoons showing the process of forming force-exerting perpendicular lateral protrusions (PLPs). Cells attached to fibers form filamentous twines that engage with with neighboring fiber (i). Subsequently, actin lamellum grows along the base of twine from cell body (ii) followed by formation of secondary twines (iii). Engagement of secondary twine with neighboring fiber leads to formation of a suspended twine-bridge (iv) that facilitates advancement of the lamellum that generates a suspended interfiber lamellum (v). Over time (~minutes), the interfiber lamellum broadens and applies contractile forces causing neighboring fiber tocontract inwards, thus creating force exerting PLPs(vii). Phase images in green and red (dashed) boxes depict two sample cases of PLPs in steps (i-vi). (c) Phase images of whole cell forming PLP. Red dashed lines indicate undeflected position of neighboring fiber. Orange arrow indicates cell migration direction. (B) Sample images of twine-bridges of varying sizes and shapes and arrows point towards the neighboring fiber.

### Actin-based development of PLPs, and dependence on fiber orientation

Filopodial extensions and their transition to lamellipodial structures are well-known to be driven by actin dynamics at the leading edge. We inquired if the twine-PLP transition was also actin-based (**Figure 4**). We recorded the growth rate of primary twines as they emerged from actin-rich ruffles and found them to extend at ~0.1 μm/s, which matches the reported kinetics of filopodia extension rates (~0.12-0.2 μm/s^45^). After engagement with neighboring fibers, the primary twines continued to grow at rates similar to-those prior to engagement (inset in **Figure 4a**). Next, we analyzed the growth of lamellae to form the twine-bridges and found that they advance at the rate of ~0.1 μm/s (**Figure 4b**), similar to actin polymerization rates (~0.12-0.14 μm/s).^39^ We saw that twine-bridges could mature into PLP structures that exert contractile force as they broadened. We measured the width of twine-bridges halfway along their span length and found them to widen at ~0.028 μm/s (**Figure 2a(vii)**), which matched the actin retrograde flow rates in the lamellipodial tip (~0.008-0.025 μm/s) but were much faster than flow rates reported in lamella (~0.004-0.008 μm/s).^46^ Overall, our measurements of twine, lamellum and twine-bridge widening rates suggested that twine-PLP transitions are similar to actin-based filopodial-lamellipodial transitions.

**FIGURE 4.**
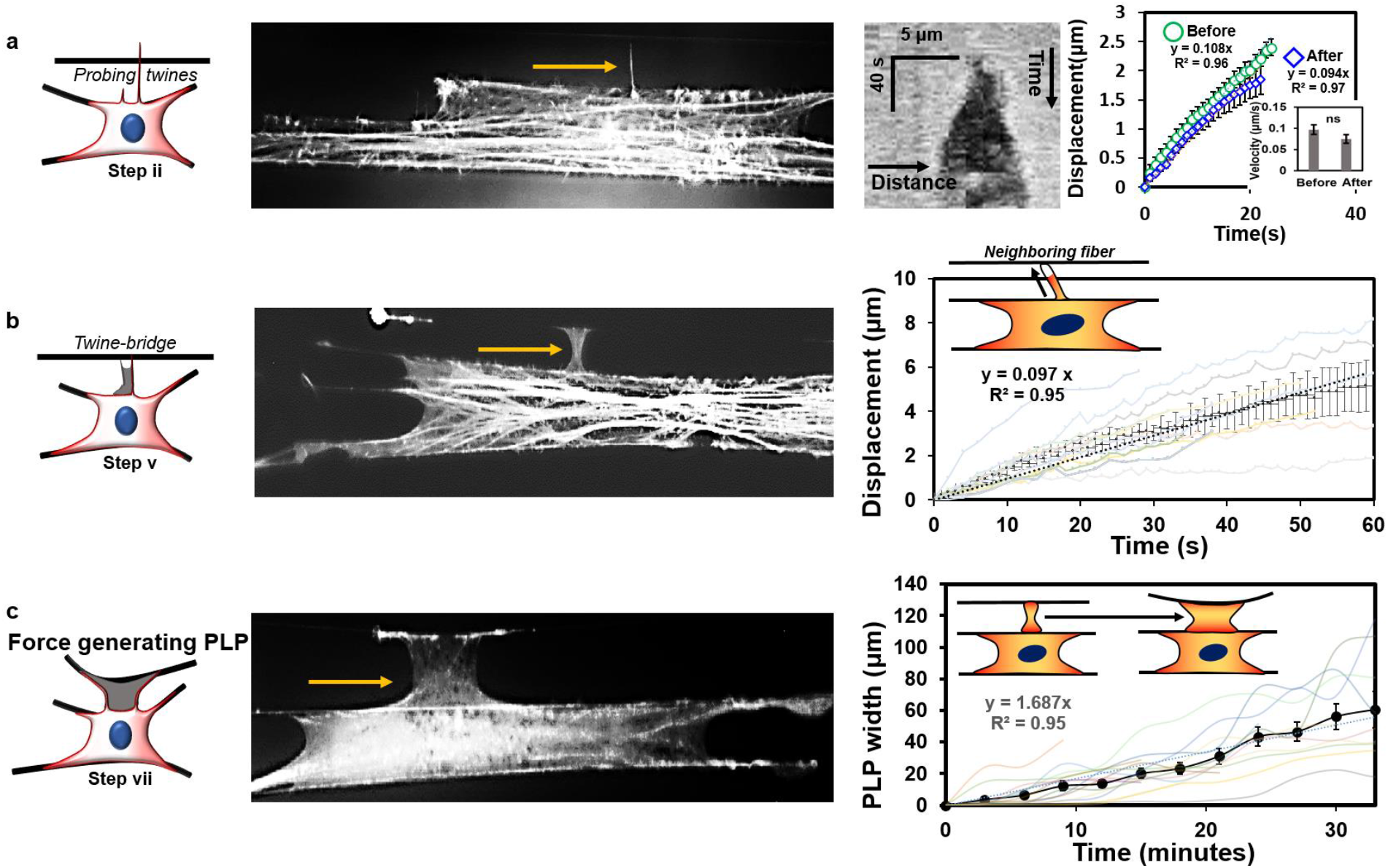
Actin kinetics in PLP formation. (a)Twines attached to neighboring fibers grow in width along the neighboring fiber axis as demonstrated by the kymograph. Kymograph was obtained by performing a line scan along the fiber axis. The average displacement vs. Time profiles show twine advancement before and after engagement with neighboring fibers (n=17 for both). Inset shows average velocity for the two categories. (b) Displacement vs. time profiles showing extension rates of actin lamellum between twine-bridges (n=13). Inset shows cartoon of actin lamellum advancement along the twine-bridge. (c) Broadening of twine-bridge into force exerting PLP (n=15). Data for linear fits to average of all profiles in included in each plot.

Next, we recorded that twines and PLPs of varying lengths (5-35 μm) formed along the entire length of the cell body (**Figure 5a**). Intriguingly, we found that twines were attached to neighboring fibers over a wide range of angles, but the ensuing PLPs were primarily oriented orthogonal to the parent cell body (**Figure 5b**). We also inquired if the spatial organization of twines and PLPs were specific to cells attached to anisotropic fiber networks. We constructed fiber networks with three more designs (hexagonal, angled, and with crosshatches) and recorded the number of twines (length ≥ 3 μm) forming in each fiber category. We found that the greatest twine formation per cell over a two-hour period (imaged every two minutes) occurred in elongated cells with high aspect ratios (observed on anisotropic and hexagonal structures, **Figure 5c**). Since twines were attaching to neighboring fibers in all categories, we calculated the number of PLPs formed from these attached twines (**Figure 5d**). Not surprisingly, we found that, on average, nearly half of engaged twines transitioned to PLPs in cells attached to anisotropic parallel fibers, whereas for all other fiber networks, we rarely encountered the formation of PLPs. Altogether, our data demonstrate that the maximum number of actin-based twines are formed in elongated cell shapes, but the transition of these twines to force-exerting PLPs almost exclusively occurs in anisotropic fiber arrangements.

**FIGURE 5.**
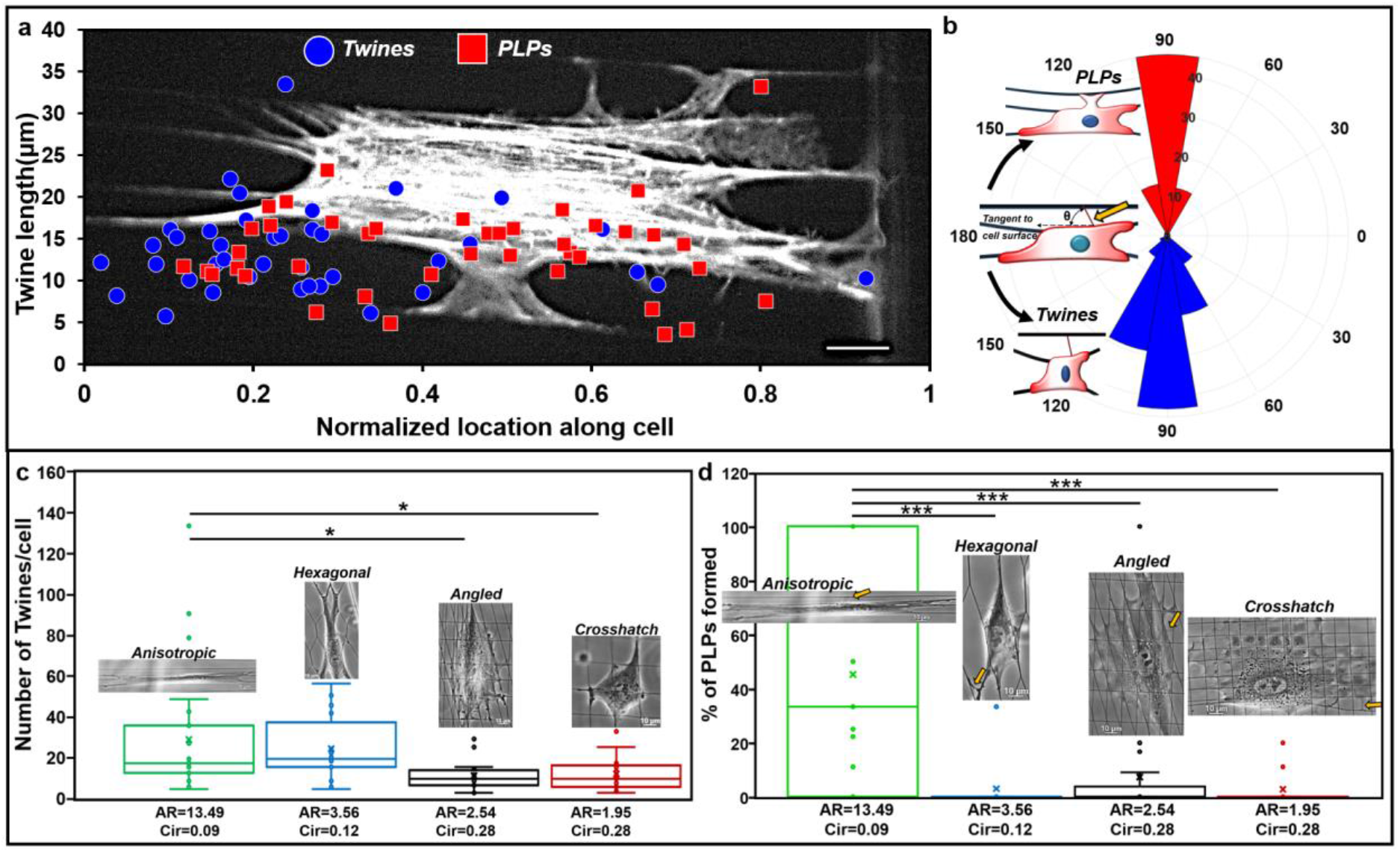
Twine and PLPs. (a) Plot showing the length of twines observed with respect to corresponding location of emergence of twines and PLPs along the normalized cell length. Backdrop shows a cell forming multiple lateral protrusions. Scale bar 10 μm. (b) Polar histogram of angle formed by twine or PLP engagement axis and a tangent to cell body (n=84 for PLP and 123 for twines). PLPs are oriented mostly perpendicularly to cell body. (c-d) Four fiber network designs were created to interrogate cell shape (AR: aspect ratio and Circ.: circularity) driven number density of (i) twines, and (ii) PLP formation ((number of PLPs/number of attached twines)*100). Elongated cells of high aspect ratios and small circularities form maximum number of twines/cell on anisotropic and hexagonal networks, while only anisotropic networks facilitate formation of PLPs. Data is analyzed from 23 randomly selected videos for each category.

### Forces of twine-bridges and PLPs

It is now well-appreciated that cells *in vivo* and in 3D interstitial ECM exert contractile forces on the neighboring fibrous environments either by pushing or tugging at individual fibers.^18,47,48^ We observed that twines that transitioned to broad PLP structures, resembling lamellipodia, were similarly able to pull neighboring fibers inwards towards the cell body, allowing cells to spread on multiple fibers. Thus, we inquired as to the development of forces in PLPs, and the resultant migratory and force response of cells as they spread from three to four and five fibers using PLPs.

First, we quantitated the forces exerted by both twines and PLPs using Nanonet Force Microscopy (NFM^32,37^, **Figure 6A**) through the establishment of tension-bearing force vectors directed towards the cell body. Specifically, the vectors originate at the paxillin attachments and point along the membrane curvature in twine-bridges and PLPs. For twine-bridges less than 6 μm in width, a single force vector pointing vertically towards the parent cell body was assigned, while two force vectors on either side of twine-bridges were assigned for sizes greater than six μm. We classified the ability of twines to exert force if they deflected the neighboring fibers ≥2 μm. Forces were estimated using a custom finite-element model that minimized the difference between experimental data (measured fiber displacement profile) and model prediction (**Figure 6a(i)**). Our analysis revealed that twines angled from the parent cell body could apply forces for only a brief period, which prevented their maturation into broad lamellipodial PLP structures (**Figure 6a(ii)**). However, we found that angled twines could transition to force exerting PLPs when the angle transitioned close to 90 degrees due to the movement/shifting of the cell body. Immunostaining for f-actin stress fibers within the lateral protrusions showed that the stress fibers in twine-bridges were oriented along the length of the twine-bridge, and those in broad PLPs were oriented along the axis of the parent cell body (yellow arrows in **Figure 6a(iii)**). Force generation increased with twine-bridge width (W, measured at the middle of span length and shown in the cartoon in **Figure 6a(iv)**), and for PLPs of widths greater than 10 μm, we observed a sharp increase in force generation, presumably due to well-defined organization of stress fibers and their alignment with the parent cell body. Overall, our force analysis indicates that transition of f-actin stress fiber orientation from orthogonal (in twines and twine-bridges) to in line with cell elongation axis (in PLPs) indicates that a neighboring fiber will become a new base fiber for further cell spreading, while inward cell contraction, pooling the extracellular fibers towards the cell body, was achieved.

**FIGURE 6.**
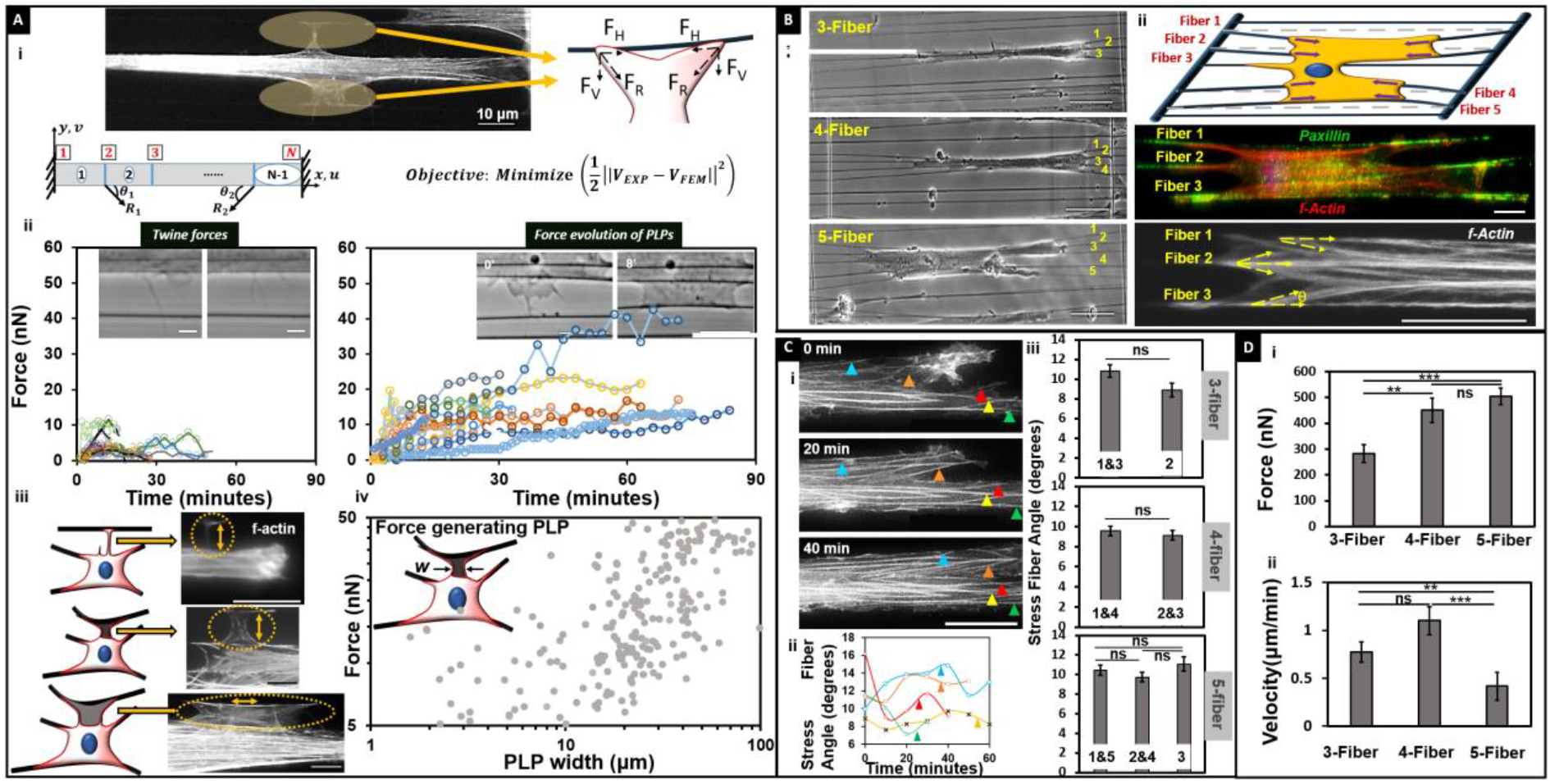
Forces applied by *t*wine-bridges facilitate cell spreading. (A) i. Representative twine-bridges (scale bar 10 μm) along with representation of forces exerted by developing twine-bridge. Force vectors are assumed to act along the membrane curvature with origin at paxillin cluster lengths. To evaluate the forces, the deflected fiber is modelled as beam with fixed-fixed boundary condition. The force vectors R_1_ and R_2_ are calculated by minimizing the objective function, where V_EXP_ represents the displacement profile of the beam measured from image sequences and V_FEM_ is the prediction from finite element model. ii. Force response of twine-bridges over time for undeveloped and developed category with inset images showing both categories. Developing twine-bridges allow cells to spread across multiple fibers. (iii) f-actin stress fibers in twines, twine-bridges and PLPs shown in dashed yellow ovals. Arrows indicate direction of f-actin stress fibers. Scale bars 10 μm. (iv) Forces in developing PLPs show an increase in force generation beyond 10 μm width (W) shown in the cartoon (n=25 PLPs). (B) i. Representative phase images of anisotropically stretched cells on 3, 4 and 5 fibers. ii. Schematic representation of NFM method to measure contractile forces of single cells attached to multiple fibers. The direction of force vectors are determined from paxillin focal adhesion clustering (FAC shown by green) acting along f-actin stress fibers (red and white). (C) i. Time-lapse images showing stress fibers tracked for angular measurements in a cell attached to five fibers. ii. Triangles of different colors in fluorescent image correspond to the transient plot. Measurements of all stress fibers could not be taken for the entire one-hour period as the respective cell front migrated out of the field of view. iii. The average f-actin stress fiber angle data: 3-fiber category: fiber 1&3 (n=53, n signifying number of stress fibers), fiber 2 (n=30). 4-fiber category: fiber 1&4 (n=80), fiber 2&3 (n=84). 5-fiber category: fiber 1&5 (n=72), fiber 2&4 (n=70), fiber 3 (n=35). (D) i. Average of total force calculated for 3 time points separated by 3 minutes each for spread cells selected randomly (n=13, 14 & 15 for 3, 4 & 5 fiber categories respectively). ii. Average migration rates for each fiber category (n=15 in each). Scale bar is (A) 20μm, (C)i. 50μm and ii. 20μm, (D)i. 20 μm.

Next, since cells were using *PLPs* in anisotropic cell migration to spread onto neighboring fibers, we inquired how the *inside-out* contractile state of cells changed when they spread from 3 to 4 and 5 fibers (**Figure 6B(i)**). For the calculation of forces applied by cells, we approximated a strategy of using a single force vector that originated at the site of paxillin focal adhesion clustering (FAC) and pointed along the dominant f-actin stress fiber (**Figure 6B(ii)**). We chose this strategy because our previous studies indicated that cells attached to ~250 nm diameter fibers formed focal adhesion clusters (FAC) at the poles.^32^ We quantified the transient change in stress fiber angles as cells migrated and observed that stress fibers maintain their relative positions (**Figure 6C(i,ii)**, **Supplementary figure S2** and **Movie M10**). Overall, stress fibers were found on average to be angled between 8-11 degrees (**Figure 6C(iii)**). The overall contractile force was computed by summation of the individual forces acting at the adhesion clusters (*F*_*cell*_ = ∑ *F*_*FAC*_). Cells spread on 5 fibers exerted higher forces (~505 nN), and as a consequence had statistically lower migration rates compared to those on 3 (~282 nN) or 4 fibers (~451 nN) (**Figure 6D**). We found that cells attached to fibers had a symmetric distribution of forces with the highest contractility driven by fibers at the outermost boundaries of cells (**Supplementary figure S3**).

## DISCUSSION and CONCLUSIONS

It has long been thought that scarring in wound healing and desmoplastic fibrotic ECM in cancer share similar origins, and newer findings point to fibroblasts as key culprits. Aligned extracellular fibers in these environments allow metastatic invasion and persistent cell migration. In contrast, the physiologically normal stroma is naturally randomly oriented and tumor suppressive^49–51^. Strategies to ablate the culprit cancer-associated fibroblasts (CAF) have indeed reduced stroma but offered no benefit to pancreatic ductal adenocarcinoma (PDAC) patients^52^, or in some cases even dangerously promoted cancer progression^53,54^. Thus, there is growing appreciation for “stroma normalization,” i.e. reinstitution of physiological ECM, and not its elimination, that is expected to impart clinical benefit^50,55–59^. A direct route to achieve stroma normalization is first understanding the biophysical links that perpetuate desmoplastic expansion, and second developing informed intervention strategies that dissuade this expansion. In this study, we describe a new functional role of anisotropic environments, which enable elongated fibroblasts to spread and contract thus continually fueling desmoplastic expansion. We show for the first time that fibroblastic cells can form contractile force-generating perpendicular lateral protrusions (PLPs) anywhere along the length of their body that contract neighboring fibers.

PLPs are formed in an orchestrated chain of events starting from actin ruffles that give rise to primary twines, which resemble microspikes and filopodia. Twines lunge and attach to neighboring fibers in a matter of seconds, which is in agreement with a recent study showing fibroblasts adhering to fibronectin in ≤ 5s in an α5β1 integrin-dependent manner.^60^ While microspikes and filopodia rarely exceed lengths of ~10 μm,^61^ the lengths of the twines observed here had a wide distribution (a short ~ 5 μm to an extremely long ~ 35 μm). To the best of our knowledge, filopodia of such long lengths have been reported only in sea urchin embryos.^62^ Filopodia in cells cultured on 2D substrates localize to the leading edge, and can transition to broad lamellipodial structures. Filopodia-lamellipodial transitions that occur at sides of cells cultured in 2D are usually associated with cell turning. In contrast, we discovered filopodia-like twines, formed in cells cultured onto anisotropic extracellular fibers, are oriented orthogonal to the parent cell, and their transition to lamellipodial PLP does not trriger a change in direction of cell migration. Instead, PLPs apply inward contractile forces that enable the cell to stretch in lateral directions, resulting in an overall elongated cell shape over an increased number of “pulled inward” fibers.

The parallel fibers used in our method mimic the aligned ECMs produced by cancer-associated fibroblasts (CAFs) *in vitro,* and the TACs_2/3_ fiber configurations *in vivo*, known to enable CAF activation^11,63^. The inter-fiber spacing (~10 μm) used in our study mimics those measured by the fibroblastic cell-derived ECM models and single harmonic generation (SHG) imaging of patient samples^7,9,63–66^. In vitro, at larger inter-fiber spacing^67,68^, cells attach to single fibers in spindle morphologies with limited instances of twine formation, akin to the behavior reported in elongated spindles in 3D gels^15^. However, the formation of PLP was similar to those observed in cells spread on multiple fibers. We also used three different fiber architectures: hexagonal, angled and crosshatched, to determine the efficacy of network designs in the formation of PLPs. Parallel and hexagonal networks support persistent cell migration in elongated shapes, while angled and crosshatched networks induce spread shapes and random migration. Elongated cell shapes on hexagonal and parallel networks form the maximum number of twines that can engage with neighboring fibers. However, only parallel fibers have the propensity to support formation of force-generating PLPs. Thus, we emphasize that ECM architecture and elongated cell shape are key physical determinants of PLP formation that is potentially needed for desmoplastic expansion.

Using parallel fibers also allowed us to use them as force sensors. While individual filopodia have been shown in the past to exert forces (~2nN)^69^, here using NFM we quantitate the transient force response of twines resembling thin to thick lamellipodia capable of exerting higher forces (tens of nano Newtons), thus providing the necessary support for cells to spread onto neighboring fibers. The increase in force generation in PLPs coincides with their increase in width (**Figure 6a(iv)**), which is correlated with the presence and spatial organization of f-actin stress fibers. Correlating the alignment of f-actin stress fibers with activation of phosphorylated focal adhesion kinase (pFAK) at adhesion sites on neighboring fibers will provide further evidence of *in vivo*-like CAF 3D adhesion formation and cells being activated to become force-generating myofibroblasts.^63^ We emphasize that our method of force measurement establishes force vectors originating from adhesion sites and directed along f-actin stress fibers, in contrast to other methods where the force vectors are determined independently of stress fiber orientation^70–72^.

In conclusion, using NFM, we show in a reproducible and replicable manner that elongated cells on anisotropic fiber networks spread laterally while contracting through the formation of force-generating lateral protrusions. Our findings identify a new cell behavior that describes the missing biophysical link in the desmoplastic expansion of aligned/anisotropic (i.e., desmoplastic) ECM. Further study of the density, organization, and size of fibers, coupled with RhoGTPase signaling in PLPs, may provide intervention strategies to deter matrix-driven fibrous spread in cancers and chronic wounding diseases.

## METHODS

### Nanonet Manufacturing and Cell Culture

The suspended polystyrene fibers were manufactured by the previously reported non-electrospinning STEP technique.^35,73^ A network of small diameter parallel fibers (~250 nm) was deposited 5-7 μm apart upon a layer of base fibers (~ 1 μm) spaced 350 μm apart and fused at the intersections. The scaffolds were placed in glass-bottom 6-well plates (MatTek, Ashland, MA) and sterilized in 70% ethanol for 10 min. After two rounds of PBS wash, 50 μl of 4μg/ml of Fibronectin (Invitrogen, Carlsbad, CA) was put on the scaffolds and incubated at 37°C for 30 min to facilitate cell adhesion. Bone Marrow-derived human Mesenchymal Stem Cells, (hMSCs; Lonza Inc, Basel, Switzerland) were cultured in supplemented growth medium (Lonza Inc) at 37°C and 5% CO2. Cells were seeded on the scaffolds by placing 50 μl droplets of 100,000 cells/mL on the suspended part, and 300 μl of media was placed around the well to prevent evaporation. After incubation for 2hr to facilitate attachment, two mL of supplemented growth medium was added to each well.

### Time-Lapse Microscopy and Cell Force Calculations

Nanonets in 6 well plates were placed in an incubating microscope (AxioObserver Z1, Carl Zeiss, Jena, Germany). Time Lapse movies were created by taking images at 20x at 3min or 40/63x at the 1-second interval with an AxioCam MRm camera (Carl Zeiss). All measurements were performed using AxioVision (Carl Zeiss) and ImageJ (National Institute of Health, Bethesda, MD). Using beam mechanics, cell forces were estimated from experimentally obtained deflection of fibers. Briefly, an optimization algorithm is written in MATLAB (MathWorks, Natick, MA) matched the experimental and computational finite-element fiber deflections to calculate forces at each time point (for details on model development, optimization algorithm and validation tests see supplementary information^33^).

### Immunohistochemistry and Immunofluorescence Imaging

Cells on fibers were fixed in 4% paraformaldehyde, permeabilized in 0.1% Triton X100 solution and blocked in 5% goat serum. Paxillin staining was done using primary rabbit anti-paxillin antibodies (Invitrogen) at a dilution of 1:250 and incubated at 4°C for 1h. Secondary goat anti-rabbit Alex Fluor 488 (Invitrogen) antibodies were incubated for 45 min at room temperature in the dark. F-Actin stress fibers were stained using Rhodamine Phalloidin. Cell nuclei were stained with 300 nM of DAPI (Invitrogen) for 5 min. The scaffolds were kept hydrated in 2ml PBS (phosphate-buffered saline) during imaging. Fluorescent images were taken using an Axio Observer Z1 microscope (Carl Zeiss). Actin live cell imaging was performed as per the manufacturer’s instructions on using the reagent CellLight Actin-RFP, Bacman 2.0 (Invitrogen).

### Statistical Analysis

Sample populations were tested for statistical significance using the Student’s t-test and analysis of variance (ANOVA) for multiple group comparisons in GraphPad software (GraphPad Prism, California). Error bars represent standard errors. Values are reported as an average ± 1 SE. * denotes p-value ≤ 0.05, ** p-value ≤ 0.01, and *** denotes p-value ≤ 0.001.

## ACKNOWLEDGMENTS

ASN, EC, and KMH are thankful to the late Professor Patricia Keely (University of Wisconsin) for discussions and guidance on cell biology and dedicate this manuscript to her. ASN would like to acknowledge the Institute of Critical Technologies and Sciences (ICTAS) at Virginia Tech for their support in conducting this study. AP and ASN would like to thank members of the STEP Lab for their helpful suggestions and discussions.

## Declaration

The authors declare no competing interests.

## Author Contributions

ASN, EC, and KMH designed research and intellectual conceptualization. ASN supervised the research. AP performed the experiments and conducted the analysis. AP, DM, and JF analyzed the IF images. KS and RKK developed the analytical and finite element model. AP, EC, KMH, and ASN wrote the manuscript.

## Funding Sources

This work is supported by National Science Foundation (1762634) awarded to ASN and KMH, by GM-R35GM122596 awarded to KMH. EC is funded by gifts donated to the memory of Judy Costin, funds from the Martin and Concetta Greenberg Pancreatic cancer Institute (Fox Chase Cancer Center), Pennsylvania’s DOH Health Research Formula Funds, the Greenfield Foundation, the 5th AHEPA Cancer Research Foundation, Inc. Fox Chase *In Vino Vita* Institutional Pilot Award, as well as NIH/NCI grants R21-CA231252 and R01-CA232256, and the Fox Chase Core grant CA06927.

## MOVIES

**<u>Movie 1</u>: PLP formation in primary human mesenchymal stem cells (hMSC)**: Force exerting perpendicular lateral protrusions form through the engagement of twines on neighboring fiber that over time mature into twine bridges. Time is shown on top left in hours:minutes:seconds

**<u>Movie 2</u>: PLPs in mouse myoblast C2C12**: Lateral protrusion (shown by arrows) that develops into PLPs. Time is shown on top left in hours:minutes: seconds

**<u>Movie 3</u>: PLPs in mouse fibroblast 3T3**: Lateral protrusion (shown by arrows) that develops into PLPs. Time is shown on top left in minutes:seconds

**<u>Movie 4:</u> PLPs in mouse embryonic fibroblast (MEF)**: Lateral protrusion (shown by arrows) that develops into PLPs. Time is shown on top left in hours:minutes:seconds

**<u>Movie 5:</u> PLPs in Human cervical tumor HeLa**: Lateral protrusion (shown by arrows) that develops into PLPs. Time is shown on top left in hours:minutes:seconds

**<u>Movie 6</u>: Actin waves along cell body**: Membrane ruffles spiral as actin wave about fiber axis. Time is shown on top left in hours:minutes:seconds

**<u>Movie 7:</u> Lateral *twines* form through membrane ruffles**: Membrane ruffles extending from cell body stratify into denser structures termed *twines.* Time is shown on top left in seconds:thousandths

**<u>Movie 8:</u> *Twine* engagement with neighboring fibers**: *Twine* engagement to neighboring fiber occurs in 1-2 s. Time is shown on top left in seconds:thousandths

**<u>Movie 9:</u> Establishment of *twine-bridges***: Growth of lamellum at the base of primary *twine* followed by the formation of secondary *twine* that facilitates establishment of primary-secondary *twine-bridge*. Time is shown on top left in seconds:thousandths

**<u>Movie 10:</u> Live f-actin stress fiber imaging**: A cell attached to five fibers is imaged for one hour to measure transient changes in f-actin stress fiber angles. The tracked stress fibers are shown by triangles next to them.

## Supplementary Figures

**Fig. S1.**
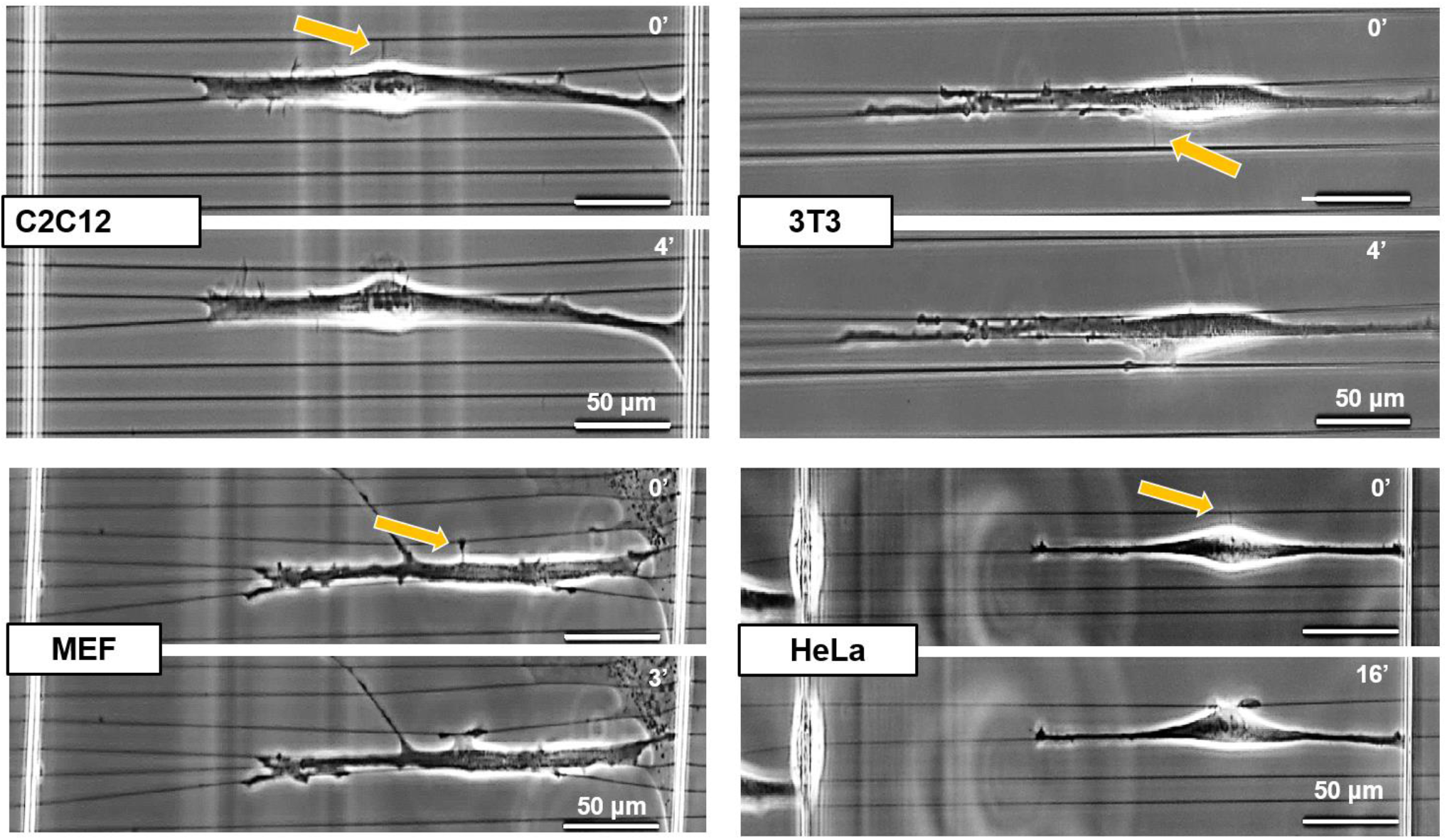
Force exerting PLPs formed by different cell lines. PLP formation through twine engagement on neighboring fibers in anisotropic environments in four different cell lines.

**Fig. S2.**
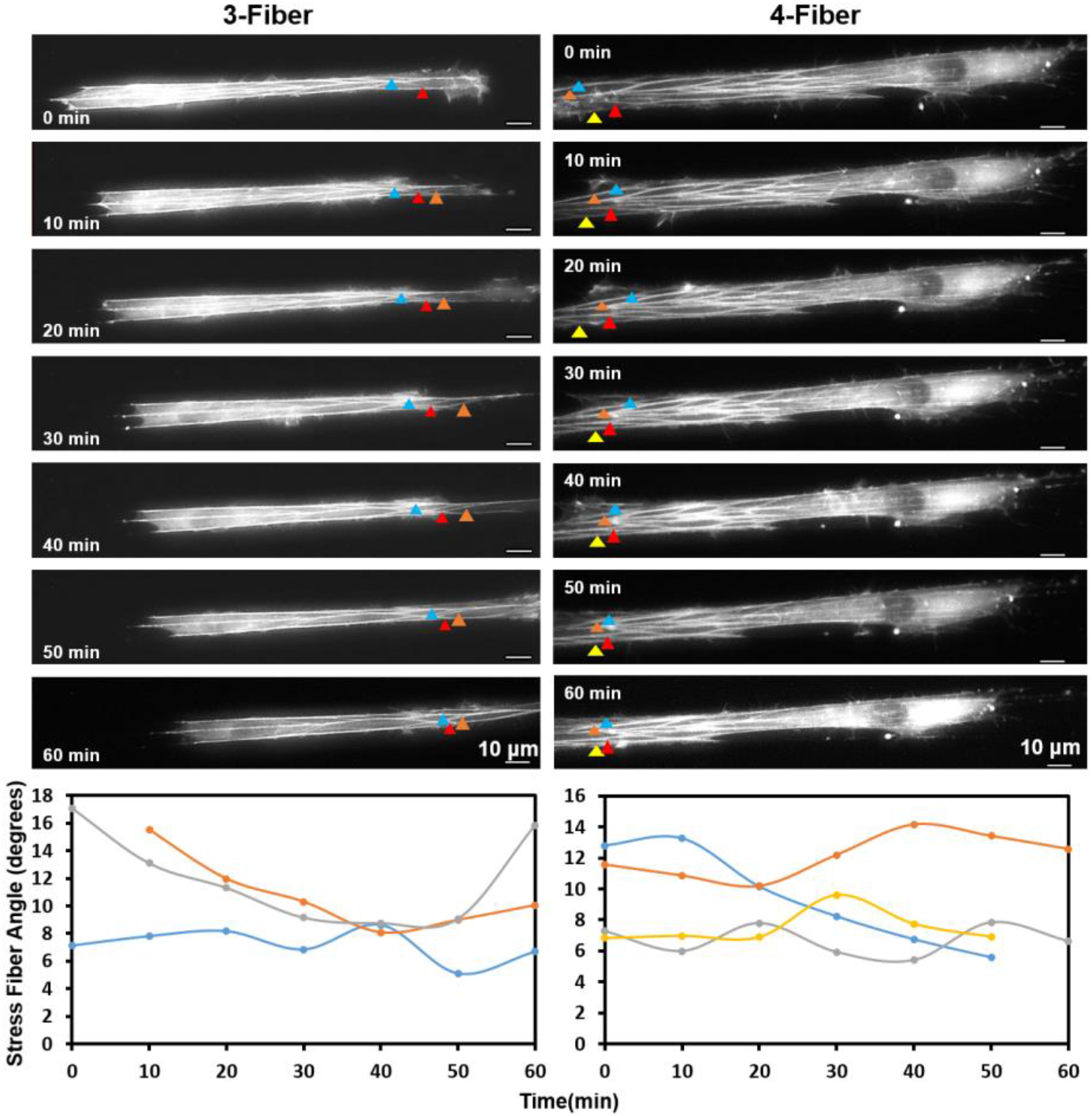
Transient analysis of f-actin stress fibers in cells attached to 3 and 4 fibers. Representative profiles of cells migrating on 3 and 4-fiber networks. Analysis shows that the stress fibers maintain their relative orientation with respect to fiber axes during cell migration.

**Fig. S3.**
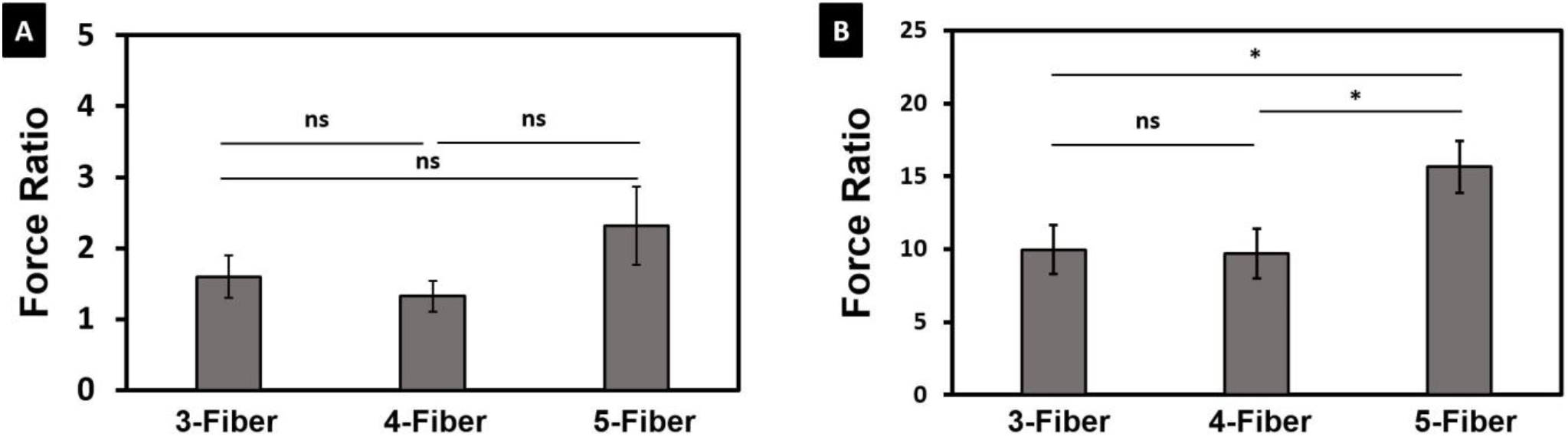
Force distribution across a cell body for cell attached to 3, 4 and 5 fibers. (A) Force ratios demonstrating symmetric distribution of forces: 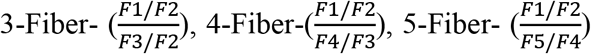 (B) Force ratios of forces at periphery to center of cell: 3-Fiber-(F1/F2), 4-Fiber-(F1/F2), 5-Fiber (F1/F3). F1 indicates force on Fiber-1, F2 force on Fiber-2, F3 force on Fiber-3, F4 force on Fiber-4, F5 force on Fiber-5. Numbering of fibers done from top to bottom (see figure 3 C(i) in main text). Sample size: n=13, 14 and 15 for 3, 4 and 5-fiber categories.

## Notes

#### Summary of Updates

New data and analysis are added.

## REFERENCES

(1) Hinz, B.; Gabbiani, G. Mechanisms of Force Generation and Transmission by Myofibroblasts. Curr. Opin. Biotechnol. 2003, 14, 538–546.

(2) Hinz, B.; Mastrangelo, D.; Iselin, C. E.; Chaponnier, C.; Gabbiani, G. Mechanical Tension Controls Granulation Tissue Contractile Activity and Myofibroblast Differentiation. Am. J. Pathol. 2001, 159, 1009–1020.

(3) Dvorak, H. F. Tumors: Wounds That Do Not Heal. Similarities between Tumor Stroma Generation and Wound Healing. N. Engl. J. Med. 1986, 315, 1650–1659.

(4) Rybinski, B.; Franco-Barraza, J.; Cukierman, E. The Wound Healing, Chronic Fibrosis, and Cancer Progression Triad. Physiol. Genomics 2014, 46, 223–244.

(5) Alexander, J.; Cukierman, E. Stromal Dynamic Reciprocity in Cancer: Intricacies of Fibroblastic-ECM Interactions. Curr. Opin. Cell Biol. 2016, 42, 80–93.

(6) Erkan, M.; Michalski, C. W.; Rieder, S.; Reiser-Erkan, C.; Abiatari, I.; Kolb, A.; Giese, N. A.; Esposito, I.; Friess, H.; Kleeff, J. The Activated Stroma Index Is a Novel and Independent Prognostic Marker in Pancreatic Ductal Adenocarcinoma. Clin. Gastroenterol. Hepatol. 2008, 6, 1155–1161.

(7) Conklin, M. W.; Eickhoff, J. C.; Riching, K. M.; Pehlke, C. A.; Eliceiri, K. W.; Provenzano, P. P.; Friedl, A.; Keely, P. J. Aligned Collagen Is a Prognostic Signature for Survival in Human Breast Carcinoma. Am. J. Pathol. 2011, 178, 1221–1232.

(8) Goetz, J. G.; Minguet, S.; Navarro-Lérida, I.; Lazcano, J. J.; Samaniego, R.; Calvo, E.; Tello, M.; Osteso-Ibáñez, T.; Pellinen, T.; Echarri, A.; et al. Biomechanical Remodeling of the Microenvironment by Stromal Caveolin-1 Favors Tumor Invasion and Metastasis. Cell 2011, 146, 148–163.

(9) Bredfeldt, J. S.; Liu, Y.; Conklin, M. W.; Keely, P. J.; Mackie, T. R.; Eliceiri, K. W. Automated Quantification of Aligned Collagen for Human Breast Carcinoma Prognosis. J. Pathol. Inform. 2014, 5, 28.

(10) Conklin, M. W.; Gangnon, R. E.; Sprague, B. L.; Van Gemert, L.; Hampton, J. M.; Eliceiri, K. W.; Bredfeldt, J. S.; Liu, Y.; Surachaicharn, N.; Newcomb, P. A.; et al. Collagen Alignment as a Predictor of Recurrence after Ductal Carcinoma In Situ. Cancer Epidemiol. Biomarkers Prev. 2018, 27, 138–145.

(11) Amatangelo, M. D.; Bassi, D. E.; Klein-Szanto, A. J. P.; Cukierman, E. Stroma-Derived Three-Dimensional Matrices Are Necessary and Sufficient to Promote Desmoplastic Differentiation of Normal Fibroblasts. Am. J. Pathol. 2005, 167, 475–488.

(12) Provenzano, P. P.; Eliceiri, K. W.; Campbell, J. M.; Inman, D. R.; White, J. G.; Keely, P. J. Collagen Reorganization at the Tumor-Stromal Interface Facilitates Local Invasion. BMC Med. 2006, 4, 38.

(13) Provenzano, P. P.; Inman, D. R.; Eliceiri, K. W.; Trier, S. M.; Keely, P. J. Contact Guidance Mediated Three-Dimensional Cell Migration Is Regulated by Rho/ROCK-Dependent Matrix Reorganization. Biophys. J. 2008, 95, 5374–5384.

(14) Heck, J. N.; Ponik, S. M.; Garcia-Mendoza, M. G.; Pehlke, C. A.; Inman, D. R.; Eliceiri, K. W.; Keely, P. J. Microtubules Regulate GEF-H1 in Response to Extracellular Matrix Stiffness. Mol. Biol. Cell 2012, 23, 2583–2592.

(15) Riching, K. M.; Cox, B. L.; Salick, M. R.; Pehlke, C.; Riching, A. S.; Ponik, S. M.; Bass, B. R.; Crone, W. C.; Jiang, Y.; Weaver, A. M.; et al. 3D Collagen Alignment Limits Protrusions to Enhance Breast Cancer Cell Persistence. Biophys. J. 2014, 107, 2546–2558.

(16) Oudin, M. J.; Jonas, O.; Kosciuk, T.; Broye, L. C.; Guido, B. C.; Wyckoff, J.; Riquelme, D.; Lamar, J. M.; Asokan, S. B.; Whittaker, C.; et al. Tumor Cell-Driven Extracellular Matrix Remodeling Drives Haptotaxis during Metastatic Progression. Cancer Discov. 2016, 6, 516–531.

(17) Adams, J. C. Cell-Matrix Contact Structures. Cell. Mol. Life Sci. 2001, 58, 371–392.

(18) Wolf, K.; Friedl, P. Mapping Proteolytic Cancer Cell-Extracellular Matrix Interfaces. Clin. Exp. Metastasis 2009, 26, 289–298.

(19) Taylor, A. C.; Robbins, E. Observations on Microextensions from the Surface of Isolated Vertebrate Cells. Dev. Biol. 1963, 7, 660–673.

(20) Nourshargh, S.; Hordijk, P. L.; Sixt, M. Breaching Multiple Barriers: Leukocyte Motility through Venular Walls and the Interstitium. Nat. Rev. Mol. Cell Biol. 2010, 11, 366–378.

(21) McEver, R. P.; Zhu, C. Rolling Cell Adhesion. Annu. Rev. Cell Dev. Biol. 2010, 26, 363–396.

(22) Ramachandran, V.; Yago, T.; Epperson, T. K.; Kobzdej, M. M.; Nollert, M. U.; Cummings, R. D.; Zhu, C.; McEver, R. P. Dimerization of a Selectin and Its Ligand Stabilizes Cell Rolling and Enhances Tether Strength in Shear Flow. Proc. Natl. Acad. Sci. U. S. A. 2001, 98, 10166–10171.

(23) Bruehl, R. E.; Springer, T. A.; Bainton, D. F. Quantitation of L-Selectin Distribution on Human Leukocyte Microvilli by Immunogold Labeling and Electron Microscopy. J. Histochem. Cytochem. 1996, 44, 835–844.

(24) Karnik, R.; Hong, S.; Zhang, H.; Mei, Y.; Anderson, D. G.; Karp, J. M.; Langer, R. Nanomechanical Control of Cell Rolling in Two Dimensions through Surface Patterning of Receptors. Nano Lett. 2008, 8, 1153–1158.

(25) Tozluoğlu, M.; Tournier, A. L.; Jenkins, R. P.; Hooper, S.; Bates, P. A.; Sahai, E. Matrix Geometry Determines Optimal Cancer Cell Migration Strategy and Modulates Response to Interventions. Nat. Cell Biol. 2013, 15, 751–762.

(26) Nelson, C. M.; Bissell, M. J. Of Extracellular Matrix, Scaffolds, and Signaling: Tissue Architecture Regulates Development, Homeostasis, and Cancer. Annu. Rev. Cell Dev. Biol. 2006, 22, 287–309.

(27) Petrie, R. J.; Doyle, A. D.; Yamada, K. M. Random versus Directionally Persistent Cell Migration. Nat. Rev. Mol. Cell Biol. 2009, 10, 538–549.

(28) Worthylake, R. a; Burridge, K. RhoA and ROCK Promote Migration by Limiting Membrane Protrusions. J. Biol. Chem. 2003, 278, 13578–13584.

(29) Pankov, R.; Endo, Y.; Even-Ram, S.; Araki, M.; Clark, K.; Cukierman, E.; Matsumoto, K.; Yamada, K. M. A Rac Switch Regulates Random versus Directionally Persistent Cell Migration. J. Cell Biol. 2005, 170, 793–802.

(30) Koons, B.; Sharma, P.; Ye, Z.; Mukherjee, A.; Lee, M. H.; Wirtz, D.; Behkam, B.; Nain, A. S. Cancer Protrusions on a Tightrope: Nanofiber Curvature Contrast Quantitates Single Protrusion Dynamics. ACS Nano 2017, 11.

(31) Mukherjee, A.; Behkam, B.; Nain, A. S. Cancer Cells Sense Fibers by Coiling on Them in a Curvature-Dependent Manner. iScience 2019, 19, 905–915.

(32) Sheets, K.; Wang, J.; Zhao, W.; Kapania, R.; Nain, A. S. Nanonet Force Microscopy for Measuring Cell Forces. Biophys. J. 2016, 111, 197–207.

(33) Tu-Sekine, B.; Padhi, A.; Jin, S.; Kalyan, S.; Singh, K.; Apperson, M.; Kapania, R.; Hur, S. C.; Nain, A.; Kim, S. F. Inositol Polyphosphate Multikinase Is a Metformin Target That Regulates Cell Migration. FASEB J. 2019, fj.201900717RR.

(34) Nain, A. S.; Sitti, M.; Jacobson, A.; Kowalewski, T.; Amon, C. Dry Spinning Based Spinneret Based Tunable Engineered Parameters (STEP) Technique for Controlled and Aligned Deposition of Polymeric Nanofibers. Macromol. Rapid Commun. 2009, 30.

(35) Wang, J.; Nain, A. S. Suspended Micro/Nanofiber Hierarchical Biological Scaffolds Fabricated Using Non-Electrospinning STEP Technique. Langmuir 2014, 30, 13641–13649.

(36) Nain, A. S.; Phillippi, J. a; Sitti, M.; Mackrell, J.; Campbell, P. G.; Amon, C. Control of Cell Behavior by Aligned Micro/Nanofibrous Biomaterial Scaffolds Fabricated by Spinneret-Based Tunable Engineered Parameters (STEP) Technique. Small 2008, 4, 1153–1159.

(37) Hall, A.; Chan, P.; Sheets, K.; Apperson, M.; Delaughter, C.; Gleason, T. G.; Phillippi, J. A.; Nain, A. Nanonet Force Microscopy for Measuring Forces in Single Smooth Muscle Cells of the Human Aorta. Mol. Biol. Cell 2017, 28, 1894–1900.

(38) Terrier, C. G.; Monzo, P.; Zhu, J.; Long, H.; Venkatraman, L.; Zhou, Y.; Wang, P.; Chew, S. Y.; Mogilner, A.; Ladoux, B.; et al. Protrusive Waves Guide 3D Cell Migration along Nanofibers. 2015, 211.

(39) Giannone, G.; Dubin-Thaler, B. J.; Rossier, O.; Cai, Y.; Chaga, O.; Jiang, G.; Beaver, W.; D??bereiner, H. G.; Freund, Y.; Borisy, G.; et al. Lamellipodial Actin Mechanically Links Myosin Activity with Adhesion-Site Formation. Cell 2007, 128, 561–575.

(40) Pontes, B.; Monzo, P.; Gole, L.; Le Roux, A.-L.; Kosmalska, A. J.; Tam, Z. Y.; Luo, W.; Kan, S.; Viasnoff, V.; Roca-Cusachs, P.; et al. Membrane Tension Controls Adhesion Positioning at the Leading Edge of Cells. J. Cell Biol. 2017, 216, 2959–2977.

(41) Zidovska, A.; Sackmann, E. On the Mechanical Stabilization of Filopodia. Biophys. J. 2011, 100, 1428–1437.

(42) Tamada, A.; Kawase, S.; Murakami, F.; Kamiguchi, H. Autonomous Right-Screw Rotation of Growth Cone Filopodia Drives Neurite Turning. J. Cell Biol. 2010, 188, 429–441.

(43) Albrecht-Buehler, G. Filopodia of Spreading 3T3 Cells: Do They Have a Substrate-Exploring Function? J. Cell Biol. 1976, 69, 275–286.

(44) Bornschlögl, T. How Filopodia Pull: What We Know about the Mechanics and Dynamics of Filopodia. Cytoskeleton 2013, 70, 590–603.

(45) Mogilner, A.; Rubinstein, B. The Physics of Filopodial Protrusion. Biophys. J. 2005, 89, 782–795.

(46) Tomasello, M.; Kirby, S.; Polich, L.; Senghas, R. J.; Wanner, E.; Gleitman, L. R.; Morford, J. P.; Coppola, M.; Kegl, J.; Senghas, A.; et al. Two Distinct Actin Networks Drive the Protrusion Of. Science (80-.). 2004, 305, 1782–1787.

(47) Hodor, P. G.; Illies, M. R.; Broadley, S.; Ettensohn, C. A. Cell-Substrate Interactions during Sea Urchin Gastrulation: Migrating Primary Mesenchyme Cells Interact with and Align Extracellular Matrix Fibers That Contain ECM3, a Molecule with NG2-like and Multiple Calcium-Binding Domains. Dev. Biol. 2000, 222, 181–194.

(48) Wyckoff, J. B.; Pinner, S. E.; Gschmeissner, S.; Condeelis, J. S.; Sahai, E. ROCK- and Myosin-Dependent Matrix Deformation Enables Protease-Independent Tumor-Cell Invasion In Vivo. Curr. Biol. 2006, 16, 1515–1523.

(49) Rybinski, B.; Franco-Barraza, J.; Cukierman, E. The Wound Healing, Chronic Fibrosis, and Cancer Progression Triad. Physiological Genomics, 2014, 46, 223–244.

(50) Schnittert, J.; Bansal, R.; Prakash, J. Targeting Pancreatic Stellate Cells in Cancer. Trends in cancer 2019, 5, 128–142.

(51) Park, C. C.; Bissell, M. J.; Barcellos-Hoff, M. H. The Influence of the Microenvironment on the Malignant Phenotype. Molecular Medicine Today, 2000, 6, 324–329.

(52) Kim, E. J.; Sahai, V.; Abel, E. V; Griffith, K. A.; Greenson, J. K.; Takebe, N.; Khan, G. N.; Blau, J. L.; Craig, R.; Balis, U. G.; et al. Pilot Clinical Trial of Hedgehog Pathway Inhibitor GDC-0449 (Vismodegib) in Combination with Gemcitabine in Patients with Metastatic Pancreatic Adenocarcinoma. Clin. Cancer Res. 2014, 20, 5937–5945.

(53) Cannon, A.; Thompson, C.; Hall, B. R.; Jain, M.; Kumar, S.; Batra, S. K. Desmoplasia in Pancreatic Ductal Adenocarcinoma: Insight into Pathological Function and Therapeutic Potential. Genes Cancer 2018, 9, 78–86.

(54) Hirata, E.; Sahai, E. Tumor Microenvironment and Differential Responses to Therapy. Cold Spring Harb. Perspect. Med. 2017, 7, 1–14.

(55) Whittle, M. C.; Hingorani, S. R. Fibroblasts in Pancreatic Ductal Adenocarcinoma: Biological Mechanisms and Therapeutic Targets. Gastroenterology 2019, 156, 2085–2096.

(56) Sherman, M. H.; Yu, R. T.; Engle, D. D.; Ding, N.; Atkins, A. R.; Tiriac, H.; Collisson, E. A.; Connor, F.; Van Dyke, T.; Kozlov, S.; et al. Vitamin D Receptor-Mediated Stromal Reprogramming Suppresses Pancreatitis and Enhances Pancreatic Cancer Therapy. Cell 2014, 159, 80–93.

(57) Neesse, A.; Krug, S.; Gress, T. M.; Tuveson, D. A.; Michl, P. Emerging Concepts in Pancreatic Cancer Medicine: Targeting the Tumor Stroma. Onco. Targets. Ther. 2013, 7, 33–43.

(58) Neesse, A.; Bauer, C. A.; Öhlund, D.; Lauth, M.; Buchholz, M.; Michl, P.; Tuveson, D. A.; Gress, T. M. Stromal Biology and Therapy in Pancreatic Cancer: Ready for Clinical Translation? Gut 2019, 68, 159–171.

(59) Bigelsen, S. Evidence-Based Complementary Treatment of Pancreatic Cancer: A Review of Adjunct Therapies Including Paricalcitol, Hydroxychloroquine, Intravenous Vitamin C, Statins, Metformin, Curcumin, and Aspirin. Cancer Manag. Res. 2018, 10, 2003–2018.

(60) Strohmeyer, N.; Bharadwaj, M.; Costell, M.; Fässler, R.; Müller, D. J. Fibronectin-Bound Α5β1 Integrins Sense Load and Signal to Reinforce Adhesion in Less than a Second. Nat. Mater. 2017, 16, 1262–1270.

(61) Saarikangas, J.; Zhao, H.; Pykäläinen, A.; Laurinmäki, P.; Mattila, P. K.; Kinnunen, P. K. J.; Butcher, S. J.; Lappalainen, P. Molecular Mechanisms of Membrane Deformation by I-BAR Domain Proteins. Curr. Biol. 2009, 19, 95–107.

(62) Mattila, P. K.; Lappalainen, P. Filopodia: Molecular Architecture and Cellular Functions. Nat. Rev. Mol. Cell Biol. 2008, 9, 446–454.

(63) Franco-Barraza, J.; Francescone, R.; Luong, T.; Shah, N.; Madhani, R.; Cukierman, G.; Dulaimi, E.; Devarajan, K.; Egleston, B. L.; Nicolas, E.; et al. Matrix-Regulated Integrin Αvβ5 Maintains Α5β1-Dependent Desmoplastic Traits Prognostic of Neoplastic Recurrence. Elife 2017, 6.

(64) Malik, R.; Luong, T.; Cao, X.; Han, B.; Shah, N.; Franco-Barraza, J.; Han, L.; Shenoy, V. B.; Lelkes, P. I.; Cukierman, E. Rigidity Controls Human Desmoplastic Matrix Anisotropy to Enable Pancreatic Cancer Cell Spread via Extracellular Signal-Regulated Kinase 2. Matrix Biol. 2019, 81, 50–69.

(65) Robinson, B. K.; Cortes, E.; Rice, A. J.; Sarper, M.; Del Río Hernández, A. Quantitative Analysis of 3D Extracellular Matrix Remodelling by Pancreatic Stellate Cells. Biol. Open 2016, 5, 875–882.

(66) Novotny, G. E.; Gnoth, C. Variability of Fibroblast Morphology in Vivo: A Silver Impregnation Study on Human Digital Dermis and Subcutis. J. Anat. 1991, 177, 195–207.

(67) Sheets, K.; Wunsch, S.; Ng, C.; Nain, A. S. Shape-Dependent Cell Migration and Focal Adhesion Organization on Suspended and Aligned Nanofiber Scaffolds. Acta Biomater. 2013, 9, 7169–7177.

(68) Sharma, P.; Ng, C.; Jana, A.; Padhi, A.; Szymanski, P.; Lee, J. S. H.; Behkam, B.; Nain, A. S. Aligned Fibers Direct Collective Cell Migration to Engineer Closing and Nonclosing Wound Gaps. Mol. Biol. Cell 2017, 28, 2579–2588.

(69) Bridgman, P. C.; Dailey, M. E. The Organization of Myosin and Actin in Rapid Frozen Nerve Growth Cones. J. Cell Biol. 1989, 108, 95–109.

(70) Beningo, K. A.; Wang, Y. L. Flexible Substrata for the Detection of Cellular Traction Forces. Trends Cell Biol. 2002, 12, 79–84.

(71) Steinwachs, J.; Metzner, C.; Skodzek, K.; Lang, N.; Thievessen, I.; Mark, C.; Münster, S.; Aifantis, K. E.; Fabry, B. Three-Dimensional Force Microscopy of Cells in Biopolymer Networks. Nat. Methods 2016, 13, 171–176.

(72) Cesa, C. M.; Kirchgessner, N.; Mayer, D.; Schwarz, U. S.; Hoffmann, B.; Merkel, R. Micropatterned Silicone Elastomer Substrates for High Resolution Analysis of Cellular Force Patterns. Rev. Sci. Instrum. 2007, 78, 034301.

(73) Jana, A.; Nookaew, I.; Singh, J.; Behkam, B.; Franco, A. T.; Nain, A. S. Crosshatch Nanofiber Networks of Tunable Interfiber Spacing Induce Plasticity in Cell Migration and Cytoskeletal Response. FASEB J. 2019, fj.201900131R.

